# A defined microbial community remodels host metabolism and pathogen resistance in *Caenorhabditis elegans*

**DOI:** 10.64898/2026.09.06.749774

**Authors:** Nathan Dennis, Laura Freeman, Mireya Vazquez-Prada, Imogen Langley, Madison S. Mortensen, Antonis A. Karamelegos, Jennifer L. Watts, Marina Ezcurra

**Author notes:** Correspondence: Marina Ezcurra, University of Kent, School of Natural Sciences, Canterbury CT2 7NZ.

## Abstract

The microbiome is an important regulator of metabolism and health, yet the mechanisms by which microbial communities influence host physiological state remain incompletely understood. Here we used a defined host-microbiota model consisting of *Caenorhabditis elegans* and an 11-member bacterial consortium representative of its natural microbiota (DefNatMta). We investigated how this native microbial community influences host metabolism and physiology relative to the standard laboratory diet, *E. coli* OP50. Transcriptomic analyses revealed remodelling of host pathways associated with lipid metabolism, immunity and xenobiotic detoxification. Comparison with fasting-responsive transcriptional programmes showed that the DefNatMta response is distinct from fasting or caloric restriction. DefNatMta enhanced resistance to *Staphylococcus aureus* infection, and this protective effect was consistent with roles for the conserved host defence regulators PMK-1/p38 MAPK and HLH-30/TFEB. Additionally, DefNatMta reduced lipid accumulation, modified expression of lipid metabolism genes, and altered fatty acid composition, including enrichment of monomethyl branched-chain fatty acids and polyunsaturated fatty acids. Together, these findings demonstrate that a defined microbial community can induce transcriptional, metabolic and physiological remodelling in the host. Our results are consistent with microbial communities shaping host metabolic state and immune competence.

## Introduction

The microbiota is a major regulator of host physiology, influencing metabolism and immunity through the production and modification of bioactive molecules. While many studies have focused on water-soluble microbial metabolites, increasing evidence suggests that microbiota-derived lipids also play important roles in host physiology. Microbial lipids can influence fatty acid synthesis, energy metabolism and immune signalling, processes that are central to how organisms acquire, store and utilise energy and respond to environmental challenges. However, how microbial communities shape host metabolism and physiological state is not well understood (Koh and Bäckhed, 2020; Lamichhane et al., 2021; Brown et al., 2023).

The nematode *Caenorhabditis elegans* provides a powerful model for studying host-microbiome interactions. Although laboratory *C. elegans* are typically maintained on monocultures of *Escherichia coli* OP50, wild animals harbour diverse native microbiota with diverse metabolic capacities that can influence host physiology (Dirksen et al., 2016; Zimmermann et al., 2020). Previous studies have shown that individual native microbiota members can alter host metabolism, immunity and stress responses. For example, *Ochrobactrum* and *Pseudomonas* isolates induce broad transcriptional changes associated with metabolism and host defence (Yang et al., 2019; Pees et al., 2024). More recently, transcriptomic analyses of animals colonised with the defined native microbial community CeMbio demonstrated that microbial communities can also induce extensive host transcriptional remodelling (Kim et al., 2025). However, how native microbial communities influence host physiology compared with conventional laboratory bacterial diets, and the functional consequences of these changes, remain largely unexplored.

To address this question, we used DefNatMta, a defined 11-member bacterial consortium designed to capture the natural taxonomic diversity of the *C. elegans* microbiota (Table 1) (Dirksen et al., 2016; Dennis et al., 2025). This consortium colonises the *C. elegans* intestine and supports normal growth and reproduction, providing a tractable model for studying the physiological effects of native microbial communities. In previous work, we showed that DefNatMta alters muscle function, mitochondrial network dynamics and ATP levels in muscle, implicating the microbiota in host metabolic regulation (Dennis et al., 2025). Building on these findings, here we investigate how DefNatMta influences host metabolism and immunity using transcriptomic, fatty acid composition and physiological analyses. We show that DefNatMta induces transcriptional and physiological remodelling distinct from fasting or caloric restriction, characterised by altered fatty acid composition, reduced lipid accumulation, and increased pathogen resistance.

**Table 1.** Composition of the DefNatMta consortium. All strains were originally isolated by Dirksen et al. (2016).

| Strain | Phylum |
| --- | --- |
| Achromobacter MYb9 | Proteobacteria |
| Acinetobacter MYb10 | Proteobacteria |
| Pseudomonas MYb11 | Proteobacteria |
| Arthrobacter MYb27 | Actinobacteria |
| Microbacterium MYb45 | Actinobacteria |
| Bacillus MYb56 | Firmicutes |
| Stenotrophomonas MYb57 | Proteobacteria |
| Ochrobactrum MYb71 | Proteobacteria |
| Leuconostoc MYb83 | Firmicutes |
| Chryseobacterium MYb120 | Bacteroidetes |
| Pseudomonas MYb218 | Proteobacteria |

## Materials and methods

### *C. elegans* Strains and Maintenance

Bristol N2 (wild type), AU78 *agIs219 [T24B8.5p::GFP::unc-54 3’ UTR + ttx-3p::GFP::unc-54 3’ UTR]*, IG274 *frIs7 [nlp-29p::GFP + col-12p::DsRed]*, ZC936 *yxEx341 [sma-6p::gfp; unc-122p::gfp]*, KU25 *pmk-1(km25)*, JIN1375 *hlh-30(tm1978)*, DMS303 *nIs590 [fat-7p::fat-7::GFP + lin15(+)]*, PHX649 *syb649 [fat-6::GFP]*, and LIU1 *ldrIs1 [dhs-3p::dhs-3::GFP + unc-76(+)]*, were obtained from the Caenorhabditis Genetics Center (CGC, University of Minnesota) and maintained as previously described (Stiernagle, 2006). All experiments were conducted at 20°C. Age-synchronised *C. elegans* populations were obtained via sodium hypochlorite-sodium hydroxide treatment as previously described (Stiernagle, 2006); egg solutions were transferred directly onto seeded NGM plates and manually transferred to fresh media at the L4 stage.

### Bacterial Strains and Growth Conditions

*E. coli* OP50 was obtained from the Caenorhabditis Genetics Center (CGC, University of Minnesota) and grown at 37°C in a 180 RPM shaking incubator for at least 12 hours. DefNatMta strains *Achromobacter sp.* F32 MYb9, *Acinetobacter sp.* LB BR12338 MYb10, *Pseudomonas lurida* MYb11, *Arthrobacter aurescens* MYb27, *Microbacterium oxydans* MYb45, *Bacillus sp.* SG20 MYb56, *Stenotrophomonas sp.* R-41388 MYb57, *Ochrobactrum sp.* R-26465 MYb71, *Leuconostoc pseudomesenteroides* MYb83, *Chryseobacterium sp.* CHNTR56 MYb120 and *Pseudomonas tuomuerensis* MYb218 (Dirksen et al., 2016) were grown at 25°C in a stationary incubator for three days before being mixed in equal volumes to form the DefNatMta consortium (Dennis et al., 2025). 200 µL aliquots of culture (*E. coli* OP50) or culture mixture (DefNatMta) were seeded onto 60 mm NGM plates and allowed to dry at room temperature for three days prior to use.

### Transcriptomic analysis via RNA-Seq

300-600 synchronised animals were collected in RNase-free centrifuge tubes and washed three times with M9. Samples were re-suspended in 500 µL TRIzol^TM^ (Invitrogen) and subjected to 10 freeze-thaw cycles at-80 °C / 37 °C, with each cycle followed by a 30 second vortex. Samples were treated with 125 µL bromo-3-chloropropane and centrifuged at 13700 x g for 10 minutes at 4 ^◦^C. The clear upper layers were transferred to fresh RNase-free centrifuge tubes and RNA extracted using an RNeasy RNA Extraction Kit (QIAGEN) in accordance with the manufacturer’s instructions. RNA eluates were collected with 35 µL of RNAse free water and stored at-20 °C.

The RNA was processed by Glasgow Polyomics. RNA libraries were prepared using the Illumina Reverse Stranded mRNA library preparation method with poly(A) selection and sequenced on an Illumina TruSeq 2000 sequencer with paired-end reads of 75 bp, averaging over 30 million reads per sample.

Data was processed using Galaxy V. 24.1.0. Adapter trimming and filtering of low quality were performed with fastp (Chen et al., 2018). Trimmed reads were mapped to the *C. elegans* genome (WBCel235) using RNA STAR with default settings (Dobin et al., 2012). Read counts were extracted using featureCounts with default settings (Liao et al., 2014). Differential expression analysis was performed using DESeq2 v. 1.48.1 (Love et al., 2014), comparing DefNatMta-treated animals with age-matched *E. coli* OP50-treated controls. Genes with log_2_ fold-changes (LFCs) *<*-0.6 or *>* 0.6 and FDR-adjusted p-values *<* 0.05 were classified as differentially expressed genes (DEGs).

### Gene Set Enrichment Analysis

Gene set enrichment analysis (GSEA) was performed using WormCat and GeneModules. WormCat was used to identify enriched functional categories among differentially expressed genes (Holdorf et al., 2020). Statistical significance was assessed using Fisher’s exact tests with Bonferroni correction for multiple comparisons, and categories with adjusted p-values < 0.05 were considered statistically significant.

GeneModules was used to analyse the activity of transcriptionally co-regulated gene modules (Cary et al., 2020). GeneModules tool 3 (module annotations) was used to identify modules for which the perturbation most highly activating the module was a form of caloric restriction, and GeneModules tool 4 was used to compare the activity of these modules in DefNatMta-treated animals relative to animals subjected to a 9-hour fast (Uno et al., 2013; GEO ID: GSE27677).

### *S. aureus* infection assay

Synchronised L4 stage animals were washed twice with M9 buffer and transferred onto unseeded NGM plates. Subsequently, 140-300 animals were transferred onto three unseeded plates to remove surface bacteria, and 70-100 animals were transferred to each TSA plate seeded with *S. aureus* or *E. coli* OP50. Animals were transferred to fresh media daily, and the number of live, dead and censored worms was recorded.

### BODIPY Staining

BODIPY staining was performed using a droplet-staining procedure adapted from a previous protocol (Klapper et al., 2011; Ezcurra et al., 2018). Briefly, groups of 15-20 animals were transferred between 20 µL droplets of M9, fixative and stain, on a 60 mm petri dish lid using an eyelash pick. Using this procedure, *C. elegans* samples were washed twice with M9 to clear bacterial debris, fixed in 4 % paraformaldehyde in PBS for 15 minutes at room temperature, subjected to three freeze/thaw cycles at-80°C / 20°C to permeabilise the cuticle, and washed three times with M9 to remove residual fixative. The samples were then incubated with a 1 µg mL^-1^ neutral lipid-specific BODIPY (493/503; Invitrogen) solution in the dark for two hours. After staining, the animals were washed three times in M9 to remove excess BODIPY prior to imaging.

### Fluorescence imaging

For reporter strain analysis (*T24B8::GFP*, *fat-6::GFP* and *fat-7::GFP*), day 1 adults were mounted on 2.5 % agarose pads and anaesthetised with 25 mM tetramisole. Fluorescence microscopy was performed using a Leica DMR compound epifluorescence microscope with a 10x objective and an Olympus Widefield Fluorescence microscope fitted with an Andor Zyla 4.2 PLUS sCMOS camera. Fixed exposure times were used. Image analysis was performed using Fiji ImageJ v. 2.15.1 (Schindelin et al., 2012) by measuring the mean grey area within the entire body of each animal and subtracting the background value of each image.

Lipid droplet diameter and quantity were assessed using the *dhs-3p::dhs-3::GFP + unc-76(+)* transgene (Na et al., 2015). Animals were mounted as described above and imaged using a Zeiss LSM880 confocal laser-scanning microscope with a 40x objective. Z-stack images were taken of the anterior intestine in the region spanning from intestinal cells int1 to int3. Z-stacks were taken over the entire range of observable fluorescence (approximately 70 µm) in 2 µm slices with a 50 % overlap between slices. BODIPY stained animals were mounted as described above and imaged using a Zeiss LSM880 confocal laser-scanning microscope with a 10x objective. Images were taken as Z-stack tile scans through the entire range of observable fluorescence (approximately 70 µm) in 6 µm slices with a 50% overlap between slices. Tile scans were automatically stitched following image acquisition using Zen Black (Zeiss). A minimum of three biological replicates were performed with a minimum of 15 animals per condition for all imaging experiments.

Image analysis for confocal images was conducted using Fiji ImageJ v. 2.15.1 (Schindelin et al., 2012). *dhs-3::GFP* and BODIPY images were maximally projected prior to analysis. To minimise the effects of variation in BODIPY uptake between staining experiments, BODIPY fluorescence was normalised using the average whole-body fluorescence of control animals. The size of *dhs-3::GFP*-tagged lipid droplets were measured manually in 25 x 25 µm cross sections drawn across the central region of the first anterior intestinal cell (int1) using ImageJ’s straight-line tool. Average lipid droplet diameter and the total number of lipid droplets were quantified for each cross section. Estimates of the total lipid droplet volume per cross section were calculated by summing the approximate volumes of all (n) lipid droplets using their measured diameters (d) and the formula for the volume of a sphere, where:

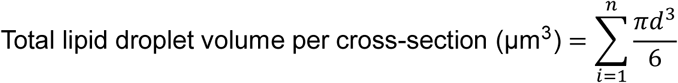

### Fatty Acid Composition Analysis

Fatty acid composition analysis was performed by gas chromatography-mass spectrometry (GC-MS) of fatty acid methyl esters (FAMEs) using a modified version of a previous protocol (Harrison and Watts, 2022). Approximately 200-1000 animals were collected into 13 x 100 mm glass tubes using warm MQ water and incubated on ice for 5 minutes. Supernatants were then aspirated and replaced with fresh MQ water, and this process was repeated until all visible bacterial contamination was removed. Samples were then incubated in 2.5 % v/v sulphuric acid in methanol for 1 h in a 70 °C water bath, cooled to room temperature and treated with 1.5 mL water mixed with 200 µl hexane to extract FAMEs. Samples were shaken vigorously and centrifuged at 12 000 x g for 1 minute. 80 µL aliquots of the upper hexane phase were removed and transferred to an SP^®^-2380 Capillary GC Column. Fatty acids were separated using an Agilent 7890 GC/5975C MS in scanning ion mode with the following program: heat to 120 ^◦^C, hold for 1 min, heat to 190 ^◦^ by 10 °C min^-1^, heat to 200 °C by 2 °C min^-1^. The relative abundance of each fatty acid was then calculated by dividing the area of each GC-MS peak by the total area of all peaks.

Bacterial samples were prepared by washing cells from three-day-old *E. coli* OP50 or DefNatMta plates (60 mm) directly into 13 x 100 mm glass tubes using warm MQ water. Bacterial samples were then pelleted by centrifugation at 5600 x g for 10 minutes and the water was carefully removed. Extraction and quantification of bacterial fatty acids was then performed as described above.

### Statistics

Statistics were calculated using R v. 4.5.0 or GraphPad Prism v. 10.1.1. Survival is presented as Kaplan-Meier curves. All other data presented represent the mean ± SEM. Analyses of transcriptomic data were performed as described in Transcriptomic analysis via RNA-Seq and Gene Set Enrichment Analysis. Differences in the relative abundance of fatty acids were assessed using Wilcoxon rank-sum tests with FDR correction for multiple comparisons. Principal component analysis of fatty acid composition data was performed with scaled relative abundances using the R function *prcomp*. Survival was analysed using the Log Rank (Mantel-Cox) test. Unless stated otherwise, all remaining data were analysed using unpaired t-tests.

## Results

### DefNatMta-mediated transcriptional alterations reflect changes in host lipid metabolism

To study the effects of the native *C. elegans* microbiota on host processes, we first performed RNA-seq analysis of young adults (day 1 of adulthood) maintained with the DefNatMta consortium, using animals maintained with *E. coli* OP50 as a control. Genes with log_2_ fold-changes (DefNatMta/OP50) greater than 0.6 (upregulated by DefNatMta) or less than-0.6 (downregulated by DefNatMta), with an FDR-corrected p-value less than 0.05, were classified as differentially expressed genes (DEGs). The DefNatMta consortium altered *C. elegans* gene expression, with 314 DEGs observed (**Figure 1A**). To determine the likely physiological impact of these alterations, we subjected each DEG list, split into upregulated and downregulated genes, to Gene Set Enrichment Analysis (GSEA) using the *C. elegans* specific web-based tool WormCat (Holdorf et al., 2020). GSEA revealed enrichment for pathogen responses, lipid metabolic processes, lipid biosynthetic processes (upregulated), detoxification/xenobiotic metabolism processes, extracellular material and transmembrane transport (downregulated; **Figure 1B**). Lipid catabolic processes (largely fatty acid β-oxidation-related genes, e.g. *ech-9*, *acox-1.5*) and pathogen responses (C-type lectins, e.g. *clec-66, clec-190*) were upregulated, and detoxification processes (cytochromes P-450 and UDP-glucuronosyltransferases, e.g. *cyp-35C1*, *ugt-41*), chromatin structure processes (e.g. *hil-1*, *hil-2*), and alternate lipid metabolic processes (largely fatty acid binding and acyl-transferase activities, e.g. *far-7*, *oac-14*) were downregulated (**Supplementary Data 1**). In summary, relative to OP50, DefNatMta induces transcriptional remodelling in *C. elegans*, notably in lipid metabolism, detoxification and immune pathways. The upregulation of β-oxidation genes and suppression of lipid-binding and acyl-transferase activities suggest altered regulation of fatty acid metabolism and storage. The upregulation of immune effectors suggests changes in response to pathogens.

**Figure 1.**
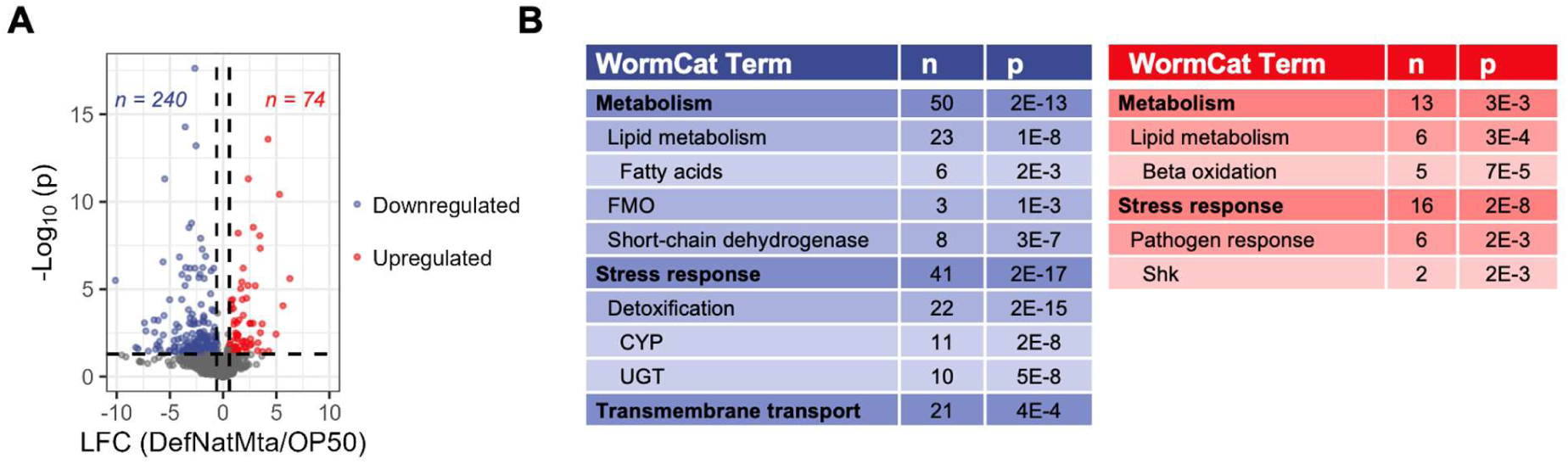
Differential gene expression induced by the DefNatMta consortium. (A) Volcano plot of gene expression of DefNatMta-fed animals relative to OP50-fed controls. Genes with an FDR-adjusted p-value of less than 0.05 and log_2_ fold-change (LFC) of less than-0.6 (downregulated; blue) or greater than 0.6 (upregulated; red) were considered differentially expressed. n=5. (B) WormCat gene set enrichment analysis of differentially expressed genes. Gene sets were split into lists of upregulated (red) and downregulated (blue) genes prior to analysis. Reported p-values represent the results of Fisher’s Exact tests with Bonferroni correction for multiple comparisons; n values represent the number of DEGs in each category.

### DefNatMta induces a transcriptional programme distinct from fasting or caloric restriction

Transcriptomic alterations in pathogen responses, lipid metabolic processes and xenobiotic metabolism are often observed during fasting and caloric restriction (McElwee et al., 2004; Chamoli et al., 2020; Wilson et al., 2021). Because DefNatMta induced transcriptional changes in these categories, we next asked whether the transcriptional response induced by DefNatMta reflects a fasting-like metabolic state. To investigate this, we compared fasting-related transcriptomic variation between DefNatMta-treated animals and animals subjected to caloric restriction using the *C. elegans*-specific GSEA tool GeneModules (**Figure 2A**). GeneModules analyses the activity of 209 groups of co-expressed genes (referred to as “modules”) derived from 1386 *C. elegans* microarray datasets and provides descriptions of each module including the top gene ontology terms, predicted transcription factor binding sites, and the perturbation (e.g. mutant vs. wild type, fasted vs. fed) most strongly activating the module. We used these descriptions to identify four modules (see **Figure 2B** Gene Ontology descriptions) for which the top perturbation was a form of caloric restriction, and compared gene expression patterns within these modules between DefNatMta-treated animals and animals subjected to a 9h fast (Uno et al., 2013; GEO ID: GSE27677). DefNatMta-treated animals showed transcriptional patterns inversely correlated with fasting across the whole transcriptome (Pearson correlation: r =-0.36, p < 0.0001) and all fasting-associated modules (Pearson correlation: all r =-0.39 to-0.57, all p < 0.0001; **Figure 2C**).

**Figure 2.**
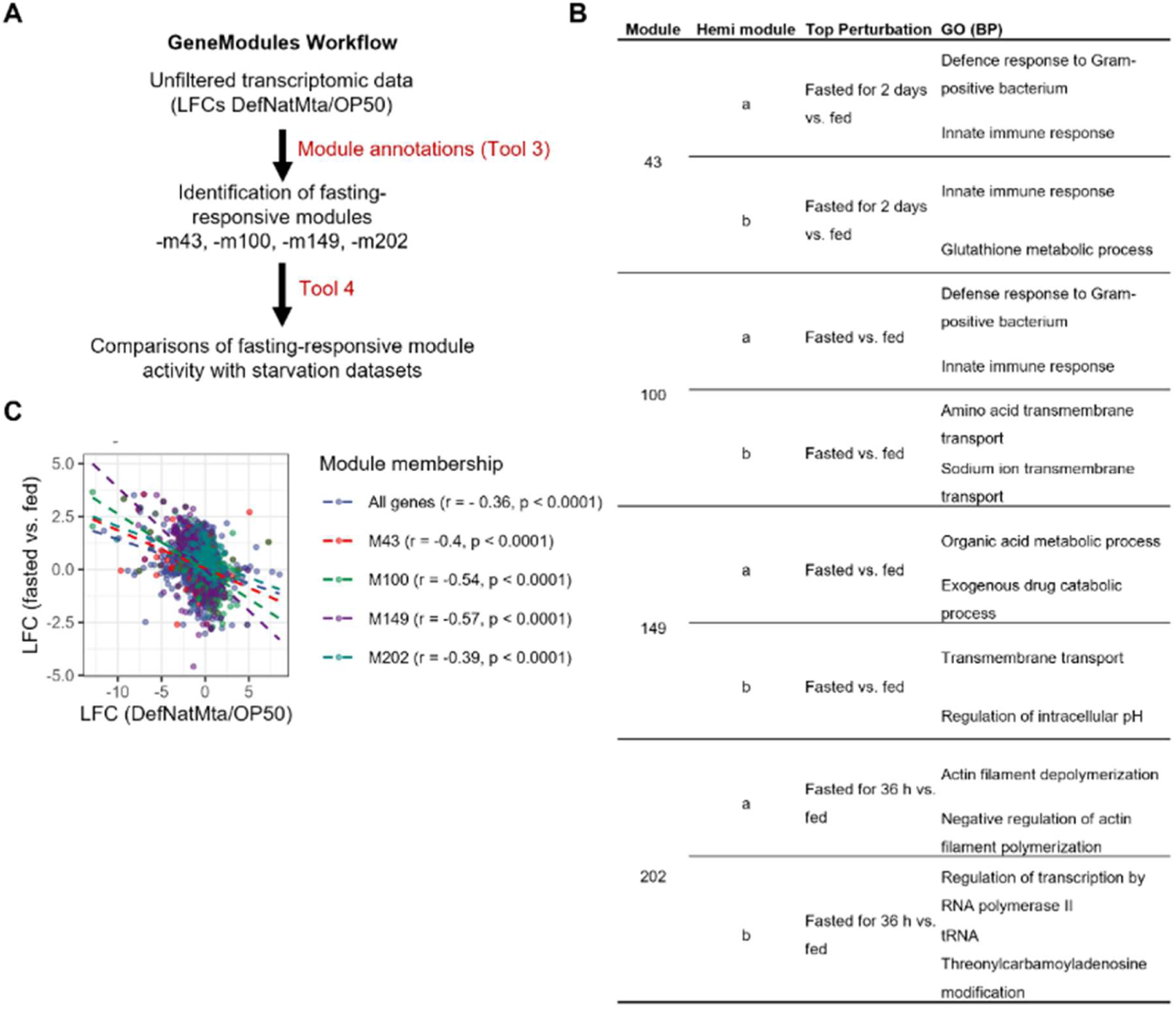
The influence of the DefNatMta consortium on fasting-responsive gene expression. (A) Description of GeneModules workflow. Module annotations (GeneModules Tool 3) were used to identify fasting responsive modules. Modules were considered fasting-responsive if the top perturbation (the condition most highly activating the module) was a form of fasting or dietary restriction. GeneModules Tool 4 was then used to compare gene expression patterns within these modules between DefNatMta-treated animals and 2-day adult animals subjected to a 9h fast. (B) Identified fasting-responsive modules, their strongest perturbation and the top two gene ontology biological process (GO BP) terms. Module descriptions were derived from the provided module annotations (GeneModules Tool 3). (C) Comparison of transcriptional variation between DefNatMta-treated animals during day 1 of adulthood, and day 2 adult worms subjected to a 9h fast. Reported r and p-values are derived from Pearson correlations. 9h starvation data is sourced from Uno et al. (2013; GEO ID: GSE27677). GeneModules tools are described in Cary et al. (2020).

This indicates that DefNatMta triggers a transcriptional programme distinct from fasting or caloric restriction, consistent with our previous work showing that *C. elegans* does not display food preference or avoidance behaviour, or alterations in developmental rate, in response to the DefNatMta consortium (Dennis et al., 2025), as might be expected if it were calorically inferior to *E. coli* OP50 (Shtonda and Avery, 2006; Stuhr and Curran, 2020). These findings suggest that the observed transcriptional changes reflect active metabolic reprogramming rather than reduced nutrient intake.

### DefNatMta remodels innate immune responses and enhances pathogen resistance

Our RNA-seq and enrichment analyses indicated that DefNatMta feeding alters host gene expression in categories associated with pathogen responses alongside metabolic pathways. In particular, genes involved in innate immune responses were significantly enriched in DefNatMta-fed worms. Differentially expressed genes included immune effectors *sysm-1, F01D5.5, skpo-1, clec-66, asp-12*, which were upregulated, and *clec-264, clec-218, clec-82, ilys-4, abf-2,* which were downregulated (**Supplementary Data 1**). In addition, expression of *sma-2* and *sma-4*, key components of the conserved DBL-1/TGF-β signalling pathway implicated in host defence against bacterial and fungal pathogens, was reduced, as was the antimicrobial peptide gene *nlp-31*. These data indicate that DefNatMta differentially regulates multiple components of the innate immune response. To validate the transcriptomic findings, we next examined fluorescent reporters representing different components of the innate immune response. We selected reporters associated with the PMK-1/p38 MAPK pathway, a central regulator of innate immunity (Troemel et al., 2006; Shivers et al., 2010; Fletcher et al., 2019), and the DBL-1/TGF-β pathway, which also contributes to host defence against bacterial and fungal pathogens (Mallo et al., 2002; Madhu et al., 2023). In addition, because antimicrobial peptides represent a major class of innate immune effectors in *C. elegans*, we examined expression of the antimicrobial peptide reporter NLP-29::GFP, as no reporter for *nlp-31* is publicly available. The *nlp-29* and *nlp-31* genes belong to the *nlp-27-nlp-31* antimicrobial peptide gene cluster, whose members are co-regulated during innate immune responses (Nathoo et al., 2001; Pujol et al., 2008). All five members of the cluster showed reduced expression following DefNatMta feeding, although only *nlp-31* reached statistical significance (**Supplementary Data 1**).

Consistent with the RNA-seq data, DefNatMta significantly increased expression of the PMK-1/p38 MAPK reporter SYSM-1::GFP by 79% (p <0.0001; **Figure 3A**). In contrast, expression of the DBL-1/TGF-β signalling reporter SMA-6::GFP was significantly reduced by 78% (p<0.001; **Figure 3B**), consistent with the downregulation of the pathway components *sma-2* and *sma-4*. Similarly, expression of the antimicrobial peptide reporter NLP-29::GFP was reduced in DefNatMta-fed animals (66%, p<0.0001; **Figure 3C**), in agreement with the reduced expression of the related antimicrobial peptide gene *nlp-31* observed in the transcriptomic analysis. Together, these findings support the conclusion that DefNatMta differentially modulates distinct innate immune pathways rather than broadly activating host immune responses.

**Figure 3.**
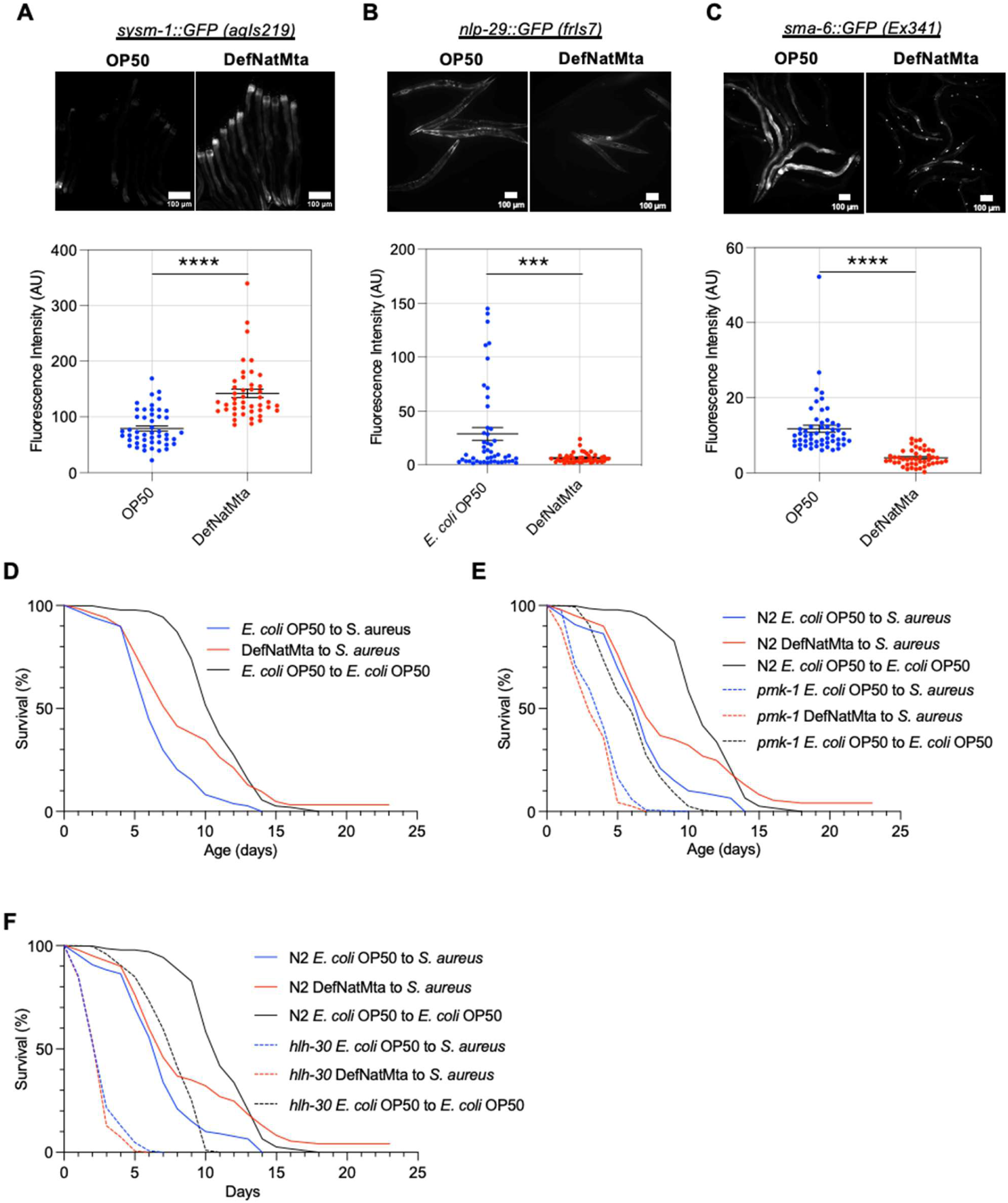
DefNatMta alters host immunity. (A) Top: Representative fluorescence images of *sysm-1::GFP* OP50-and DefNatMta-fed animals. Bottom: Quantification of *sysm-1::GFP* fluorescence intensity. n = 44-45. Scale bars, 100 µm. (B) Top: Representative fluorescence images of *sma-6::GFP* animals. Bottom: Quantification of *sma-6::GFP* fluorescence intensity. n = 46-48. Scale bars, 100 µm. (C) Top: Representative fluorescence images of *nlp-29::GFP* animals. Bottom: Quantification of *nlp-29::GFP* fluorescence intensity. N = 46-55. Scale bars, 100 µm. Data analysed using unpaired t-tests. Dots show fluorescence intensity from each animal. Lines show mean ± SEM. (D-F) Survival curves of OP50-and DefNatMta-fed animals infected with *S. aureus* at L4 stage. (D) DefNatMta increases survival in infected animals. (E) DefNatMta does not increase survival in *pmk-1* mutants. (F) DefNatMta does not increase survival in *hlh-30* mutants. ****, p < 0.0001; ***, p < 0.001.

Having established that DefNatMta induces immune-associated molecular responses, we next tested whether these molecular changes translate into altered host defence. To do this, animals were cultivated either with OP50 or DefNatMta as a bacterial food source throughout development. At the L4 stage, animals were moved to TSA plates with bacterial lawns consisting of the pathogen *Staphylococcus aureus*, and survival was recorded daily. Survival differences were assessed using Kaplan-Meier analysis and log-rank tests. Control animals were cultivated with OP50 on TSA plates throughout the experiment. We attempted to include animals cultivated with DefNatMta on TSA plates as an additional control; however, all animals cultivated with DefNatMta on TSA plates died within 24 h. As expected, exposure to *S. aureus* substantially increased risk of death compared to animals maintained on OP50 (log-rank test, P < 0.0001; HR = 3.61, 95% CI 3.01-4.23; **Figure 3D**; **Supplementary Table 1**). DefNatMta significantly improved survival following *S. aureus* infection and was associated with a reduced risk of death compared with OP50-fed controls (log-rank test, P < 0.0001; HR = 0.60, 95% CI 0.51-0.70; **Figure 3D**, **Supplementary Table 1**). As DefNatMta shortens rather than extends lifespan under standard laboratory conditions (Dennis et al., 2025), the improved survival observed does not reflect a general longevity effect. Instead, these findings are consistent with DefNatMta specifically enhancing resistance to *S. aureus* infection, indicating that the immune-associated transcriptional changes induced by DefNatMta are accompanied by improved host defence.

We next investigated whether PMK-1/p38 MAPK signalling contributes to the protective effect of DefNatMta, by examining survival of *pmk-1* mutant animals following *S. aureus* infection. As in our previous experiment, DefNatMta significantly protected against death following *S. aureus* infection compared with OP50-fed controls (log-rank test, P < 0.0001; HR = 0.68, 95% CI 0.55–0.85; **Figure 3E**, **Supplementary Table 1**). In contrast, DefNatMta did not improve survival of *pmk-1* mutants during *S. aureus* infection. Instead, DefNatMta-fed *pmk-1* mutants exhibited significantly reduced survival and an increased risk of death compared with OP50-fed *pmk-1* controls (log-rank test, P = 0.0002; HR = 1.31, 95% CI 1.10–1.57; **Figure 3E, Supplementary Table 1**). These findings are consistent with a role for PMK-1 in the protective effect of DefNatMta during pathogen challenge.

In addition to innate immune effectors, DefNatMta altered the expression of genes associated with stress responses and metabolism, suggesting that the consortium may influence host pathways beyond canonical immune signalling. We therefore examined the involvement of HLH-30, the *C. elegans* ortholog of the mammalian transcription factor TFEB, which coordinates antimicrobial, cytoprotective, and metabolic responses during infection (Visvikis et al., 2014), by challenging *hlh-30* mutants with *S. aureus*, and measuring survival. Consistent with previous studies (Visvikis et al., 2014), exposure to *S. aureus* significantly reduced survival in *hlh-30* mutants compared with OP50-fed controls (log-rank test, P < 0.0001; HR = 4.90, 95% CI 4.07–5.90; **Figure 3F**, **Supplementary Table 1**). Unlike wild-type animals, DefNatMta-fed *hlh-30* mutants did not exhibit a significant survival benefit compared with OP50-fed *hlh-30* controls (log-rank test, P = 0.063; HR = 1.11, 95% CI 0.96-1.28; **Figure 3F, Supplementary Table 1**). Together, these data suggest that DefNatMta enhances resistance to *S. aureus* through conserved host defence pathways.

### DefNatMta reduces intestinal and whole-body lipid accumulation in *C. elegans*

Having established a functional effect of DefNatMta on host defence, we next investigated the consequences of the changes in lipid metabolism identified by our transcriptomic analysis. Given that DefNatMta altered the expression of lipid metabolic genes such as fatty acid desaturases and genes associated with β-oxidation, we examined whether these transcriptional changes were accompanied by differences in lipid accumulation. *C. elegans* lacks dedicated adipose tissue and most of its lipid accumulates in lipid droplets, primarily in the intestine (Mak, 2012). We therefore used two complementary techniques to examine intestinal and whole-body stores. First, we measured lipid droplet size and quantity in the anterior intestine (**Figure 4A, B**; intestinal cell int1) on day 1 of adulthood using a transgenic strain expressing GFP-tagged DHS-3, a short-chain dehydrogenase/reductase that localises to lipid droplets and is widely used as a lipid-droplet marker (Na et al., 2015). DefNatMta-fed animals showed a pronounced reduction in lipid droplet diameter (**Figure 4C**) and a small increase in lipid droplet quantity (**Figure 4D**). Despite the higher droplet count, the total estimated lipid droplet volume per cross-section was significantly reduced in DefNatMta-fed animals (**Figure 4E**). Next, we assessed whole-body neutral lipid levels using a neutral lipid-specific BODIPY dye (BODIPY 493/503), which indicated that DefNatMta-fed animals had substantially lower neutral lipid accumulation relative to animals treated with *E. coli* OP50 (**Figure 4F, G**).

**Figure 4.**
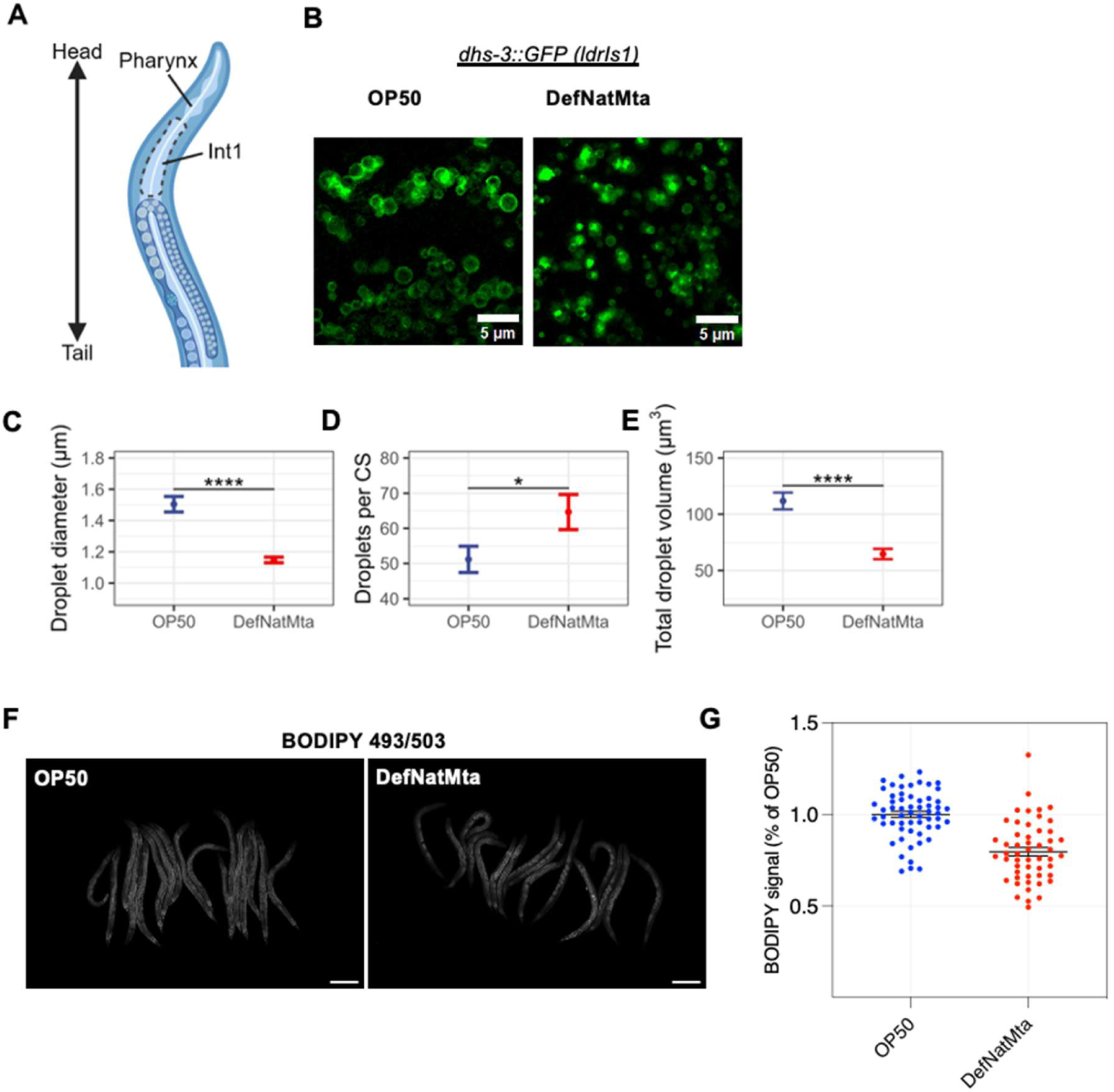
Reductions in lipid storage induced by the DefNatMta consortium. (A) Cartoon of the anterior portion of *C. elegans*, highlighting the location of intestinal cell int1. (B) Images of GFP-tagged lipid droplets in the anterior intestine (int1). The insets show representative cross-sections used for measuring lipid droplets. Scale bars, 5 µm. (C-E) Quantification of average lipid droplet diameter (C), quantity (D) and total volume (E) per cross-section. n = 11-22. (F-G) Whole-body neutral lipid storage of day 1 adults assessed via BODIPY 493/503 fluorescence. Scale bars, 200 µm. Data are presented as mean ± SEM. n = 52-58. ****, p < 0.0001; **, p < 0.01; unpaired t-tests.

### DefNatMta has a distinct fatty acid profile that is partially reflected in the host

We next investigated whether DefNatMta also alters host fatty acid composition. To distinguish differences associated with the bacterial diet from those observed in the host, we analysed the fatty acid component of DefNatMta and OP50 lawns grown on NGM plates, together with young adult *C. elegans* grown on respective bacterial diet. The two bacterial diets differed markedly in fatty acid composition. Compared with OP50, DefNatMta contained lower relative levels of saturated and cyclopropyl fatty acids and substantially higher levels of monomethyl branched-chain fatty acids (mmBCFAs), which were not detected in OP50 (**Figure 5A**, **Supplementary Data 2**). The fatty acid composition of *C. elegans* partially reflected these dietary differences, with DefNatMta-fed animals containing higher relative levels of mmBCFAs and lower levels of cyclopropyl fatty acids than OP50-fed animals (**Figure 5A**). Another notable difference between the bacterial diets was oleic acid (OA; C18:1n-9), which was abundant in DefNatMta (**Figure 5B**) but was not detected in OP50 in our analysis, consistent with previous studies (Perez and Van Gilst, 2008; Deline et al., 2013). Oleic acid has previously been shown to have beneficial effects in *C. elegans*, including enhanced resistance to infection and lifespan extension (Anderson et al., 2019; Castillo-Quan et al., 2023). In *C. elegans*, OA is also an important precursor for the synthesis of polyunsaturated fatty acids (PUFAs). DefNatMta-fed animals contained higher relative levels of PUFAs than OP50-fed animals, suggesting that the increased dietary availability of OA may contribute to this difference.

**Figure 5.**
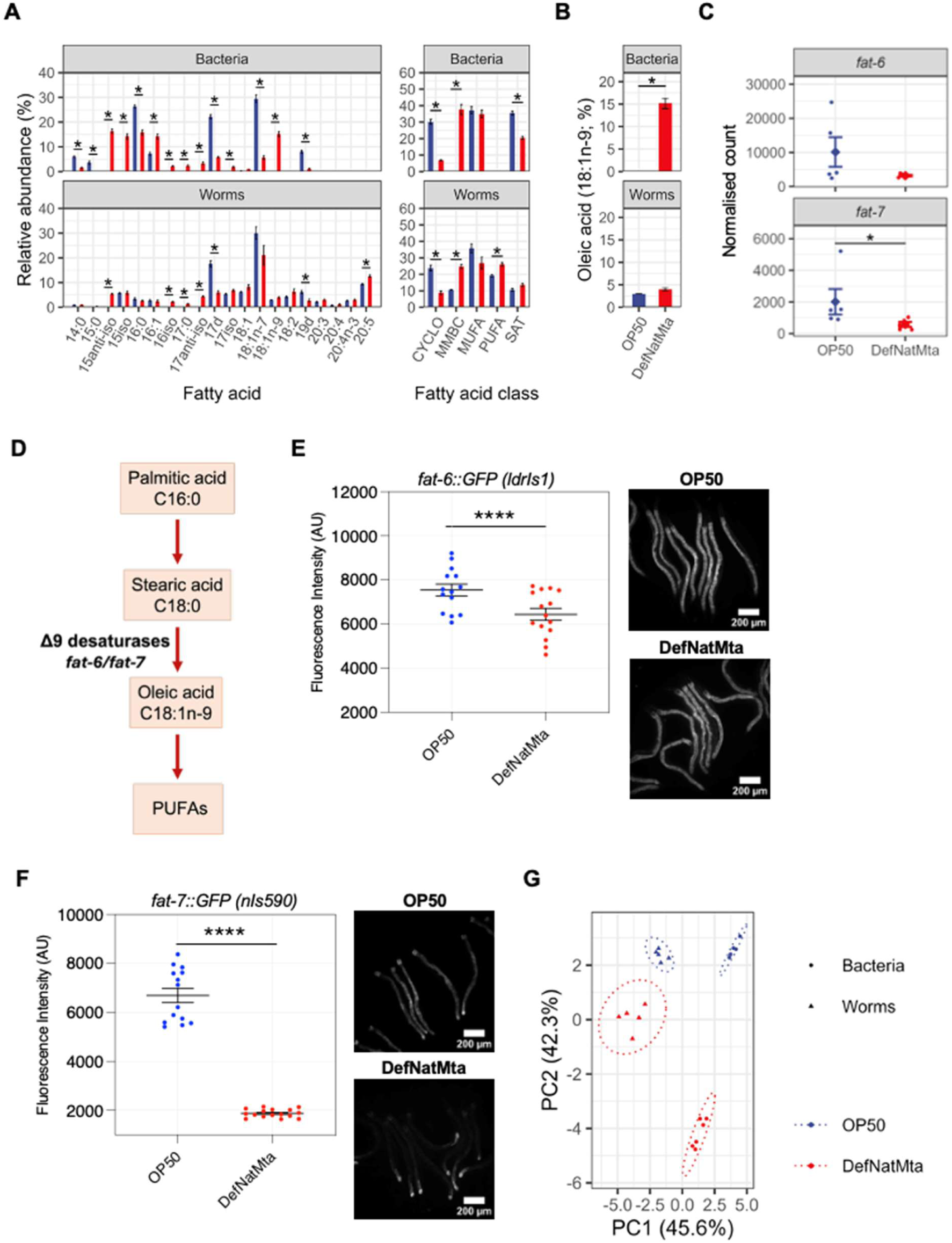
Fatty acid composition of the DefNatMta consortium and DefNatMta-fed *C. elegans*. (A) Relative abundance of individual fatty acids (left) and fatty acid classes (right) in bacterial and *C. elegans* samples comparing *E. coli* OP50 and the DefNatMta consortium. Fatty acid classes are cyclopropyl fatty acids (CYCLO), monomethyl branched-chain fatty acids (MMBC), monounsaturated fatty acids (MUFA), polyunsaturated fatty acids (PUFA) and saturated fatty acids (SAT). Bacterial fatty acids were extracted directly from three-day-old bacterial lawns on NGM plates, and *C. elegans* fatty acids were extracted from 200-1000 day-1-adult animals maintained on these lawns. n = 5. (B) Relative abundance of oleic acid (C18:1n-9) in bacterial and *C. elegans* samples maintained on OP50 or DefNatMta. (C) Normalised read counts of Δ9 fatty acid desaturases *fat-6* and *fat-7* in DefNatMta-fed animals relative to OP50 control. n = 5. p-values are derived from the differential expression model. (D) Simple overview diagram of fatty acid desaturation in *C. elegans*. FAT, fatty acid desaturase; ELO, elongase. Δ9 desaturases are highlighted blue. (E) Fluorescence intensity of individual animals expressing *fat-6::GFP* (left) and representative fluorescence images of *C. elegans* expressing *fat-6::GFP* (right). (F) Fluorescence intensity of individual animals expressing *fat-7::GFP* (left) and representative fluorescence images of *C. elegans* expressing *fat-7::GFP* (right). n = 29-30 per condition for both reporters. Scale bars, 200 µm. (G) Principal component analysis of relative fatty acid data shown in panel A. The ellipses represent the 95 % confidence intervals. n = 5. Differences in fatty acid abundance (A, B) were assessed using Wilcoxon rank-sum tests with FDR correction for multiple comparisons. FAT-6::GFP and FAT-7::GFP fluorescence (E, F) was analysed using unpaired t-tests. ****, p < 0.0001; ***, p < 0.001; **; p < 0.01; *, p < 0.05.

We next examined whether DefNatMta altered the expression of the Δ9 fatty acid desaturases involved in OA synthesis. RNA-seq analysis showed significantly reduced expression of *fat-7* in DefNatMta-fed animals, while *fat-6* showed a similar but non-significant trend (**Figure 5C**). FAT-6 and FAT-7 are Δ9 fatty acid desaturases that catalyse the synthesis of OA from stearic acid (**Figure 5D**). Consistent with the RNA-seq data, FAT-6::GFP and FAT-7::GFP translational reporters were both markedly reduced in DefNatMta-fed animals (**Figure 5E, F**). These findings are consistent with increased dietary OA reducing the requirement for endogenous OA synthesis. Principal component analysis (PCA) further demonstrated distinct fatty acid profiles across the four sample groups, with PC1 separating bacterial and *C. elegans* samples and PC2 separating DefNatMta-and OP50-associated samples (**Figure 5G**). Together, these findings show that DefNatMta differs markedly from OP50 in fatty acid composition and that these dietary differences are partially reflected in the host. DefNatMta feeding is also associated with changes in host fatty acid metabolism, including reduced expression of the oleic-acid desaturases FAT-6 and FAT-7 and increased relative levels of PUFAs.

## Discussion

Most studies of host-microbiome interactions in *C. elegans* rely on single bacterial laboratory strains, which differ substantially from the complex microbial communities encountered in nature. As a result, the extent to which native microbiotas shape host physiology and metabolism remains incompletely understood. Here, we investigated the impact of a defined native *C. elegans* microbial consortium, DefNatMta, on host physiology and metabolism. DefNatMta induced broad transcriptional remodelling in lipid metabolic, immune and detoxification pathways that was distinct from fasting or caloric restriction responses. These changes were accompanied by altered host fatty acid composition, reduced fat accumulation and enhanced pathogen resistance, showing that native microbial communities can substantially influence host physiology in *C. elegans*.

DefNatMta induced transcriptional reprogramming in *C. elegans*, with pronounced changes in genes associated with lipid metabolism, pathogen responses, and detoxification. Similar transcriptional signatures have been reported in response to individual native microbiome members such as *Ochrobactrum* MYb71 and MYb237, *Pseudomonas* MYb11 and MYb131 (Yang et al., 2019; Pees et al., 2024) as well as the defined microbial community CeMbio (Kim et al., 2025). Together, these studies suggest that modulation of metabolic, immune, and detoxification pathways represents a general feature of *C. elegans*-microbiome interactions. Our findings extend these observations by showing that a native microbial consortium can substantially remodel host lipid metabolism, fatty acid composition and lipid accumulation.

Reduced lipid accumulation, altered lipid metabolism, and enhanced stress responses are frequently observed during nutrient limitation in *C. elegans* as well as other species (Fontana and Partridge, 2015; Kapahi et al., 2016; Wilson et al., 2021), raising the possibility that DefNatMta could act as a calorically inferior food source. However, transcriptomic comparisons with fasting-associated gene modules revealed largely opposing expression patterns, arguing against a canonical fasting-like response. These findings are also consistent with our previous observations that DefNatMta does not impair development or induce food avoidance behaviours in *C. elegans* (Dennis et al., 2025). Together, these results suggest that the observed phenotypes are unlikely to arise solely from caloric restriction and instead reflect active host responses to the composition of the native microbial community.

One of the most pronounced effects of DefNatMta was the remodelling of host lipid metabolism. DefNatMta differed substantially from *E. coli* OP50 in fatty acid composition, including enrichment of monomethyl branched-chain fatty acids (mmBCFAs) and oleic acid (OA). These dietary differences were partially reflected in the host, with DefNatMta-fed animals showing increased relative levels of mmBCFAs. Notably, DefNatMta contained substantial levels of OA, which was not detected in OP50. Consistent with the increased dietary availability of OA, DefNatMta-fed animals showed reduced expression of FAT-6 and FAT-7, the Δ9 fatty acid desaturases responsible for endogenous OA synthesis. This may represent an adaptive response to increased dietary OA, reducing the requirement for endogenous OA synthesis. DefNatMta-fed animals also contained higher relative levels of PUFAs, which could potentially reflect increased availability of dietary OA as a precursor for downstream PUFA synthesis. In parallel with these changes in fatty acid composition, DefNatMta markedly reduced intestinal and whole-body lipid storage. The mechanistic relationship between altered fatty acid composition and reduced lipid accumulation remains to be determined. Our findings are consistent with previous studies showing that dietary and microbial fatty acids can influence conserved lipid-regulatory pathways in both *C. elegans* and mammals (Nomura et al., 2010; Jeon and Osborne, 2012; Kersten, 2014; Brown et al., 2023).

In addition to metabolic remodelling, DefNatMta enhanced resistance to *S. aureus* infection and induced transcriptional changes in multiple innate immune effectors. Similar effects have been reported for individual members of the native *C. elegans* microbiota (Kissoyan et al., 2019; Yang et al., 2019; Zárate-Potes et al., 2022; Pees et al., 2024). The loss of DefNatMta-mediated protection in *pmk-1* and *hlh-30* mutants is consistent with roles for PMK-1/p38 MAPK and HLH-30/TFEB signalling in the enhanced pathogen resistance. Together, these observations support the idea that native microbiota actively shape host physiological state and immune competence in *C. elegans*.

Together with previous studies of native microbiota members and microbial communities, our findings suggest that metabolism, immunity and detoxification are among the host processes most consistently influenced by microbiota exposure in *C. elegans*. However, whether these responses are mechanistically linked remains unknown. DefNatMta represents only part of the native *C. elegans* microbiota and may not capture its full ecological diversity. Because survival of *pmk-1* and *hlh-30* mutants on DefNatMta was not assessed in the absence of pathogen challenge, we cannot exclude genotype-specific effects of DefNatMta on baseline survival. Furthermore, our analyses focused primarily on fatty acids and did not directly distinguish between microbial lipid incorporation and host lipid remodelling. Future work should identify the bacterial taxa and metabolites responsible for these effects and determine how microbial communities influence the physiological state of *C. elegans*.

## Author contributions

ND and ME designed the study, prepared the figures, and wrote the manuscript. ND, LF, MVP, MSM, IL and AAK performed the experiments. ND, LF, IL and MSM analysed the data. JLW designed the fatty acid composition experiments. All authors approved the submitted manuscript.

## Acknowledgements

We thank the CGC, which is funded by NIH Office of Research Infrastructure Programs (P40 OD010440), for providing *C. elegans* strains for this study. ZC936 was a kind gift from Professor Yun Zhang, Department of Organismic and Evolutionary Biology, Harvard University.

